# Genome-wide association study of 1 million people identifies 111 loci for atrial fibrillation

**DOI:** 10.1101/242149

**Authors:** Jonas B. Nielsen, Rosa B. Thorolfsdottir, Lars G. Fritsche, Wei Zhou, Morten W. Skov, Sarah E. Graham, Todd J. Herron, Shane McCarthy, Ellen M. Schmidt, Gardar Sveinbjornsson, Ida Surakka, Michael R. Mathis, Masatoshi Yamazaki, Ryan D. Crawford, Maiken E. Gabrielsen, Anne Heidi Skogholt, Oddgeir L. Holmen, Maoxuan Lin, Brooke N. Wolford, Rounak Dey, Håvard Dalen, Patrick Sulem, Jonathan H. Chung, Joshua D. Backman, David O. Arnar, Unnur Thorsteinsdottir, Aris Baras, Colm O’Dushlaine, Anders G. Holst, Xiaoquan Wen, Whitney Hornsby, Frederick E. Dewey, Michael Boehnke, Sachin Kheterpal, Seunggeun Lee, Hyun M. Kang, Hilma Holm, Jacob Kitzman, Jordan A. Shavit, José Jalife, Chad M. Brummett, Tanya M. Teslovich, David J. Carey, Daniel F. Gudbjartsson, Kari Stefansson, Goncalo R. Abecasis, Kristian Hveem, Cristen J. Willer

**Affiliations:** Department of Internal Medicine, Division of Cardiovascular Medicine, University of Michigan, Ann Arbor, Michigan, United States.; Department of Human Genetics, University of Michigan, Ann Arbor, Michigan, United States.; deCODE genetics/Amgen, Inc., Reykjavik, Iceland.; HUNT Research Centre, Department of Public Health and General Practice, Norwegian University of Science and Technology, Levanger, Norway.; K.G. Jebsen Center for Genetic Epidemiology, Department of Public Health, Norwegian University of Science and Technology, Trondheim, Norway.; Department of Computational Medicine and Bioinformatics, University of Michigan, Ann Arbor, Michigan, United States.; Laboratory of Molecular Cardiology, Department of Cardiology, The Heart Centre, Copenhagen University Hospital, Rigshospitalet, Copenhagen, Denmark.; Department of Internal Medicine, Center for Arrhythmia Research, University of Michigan, Ann Arbor, Michigan, United States.; Regeneron Genetics Center, Tarrytown, NY, United States.; Department of Biostatistics, Center for Statistical Genetics, University of Michigan, Ann Arbor, Michigan, United States.; Department of Anesthesiology, University of Michigan, Ann Arbor, Michigan; Medical Device Development and Regulation Research Center, The University of Tokyo, Japan.; Department of Cardiology, St. Olav’s University Hospital, Trondheim, Norway.; Department of Medicine, Levanger Hospital, Nord-Trøndelag Hospital Trust, Levanger, Norway.; Department of Circulation and Medical Imaging, Norwegian University of Science and Technology, Trondheim, Norway.; Department of Cardiology, St. Olav’s University Hospital, Trondheim University Hospital, Norway.; Faculty of Medicine, University of Iceland, Reykjavik, Iceland; Department of Medicine, Landspitali - National University Hospital, Reykjavik, Iceland; School of Engineering and Natural Sciences, University of Iceland, Reykjavik, Iceland.; Department of Pediatrics and Communicable Diseases, University of Michigan, Ann Arbor, MI; Fundación Centro Nacional de Investigaciones Cardiovasculares (CNIC), Madrid and CIBERCV, Spain.; Geisinger Health System, Danville, PA 17822, USA.

## Abstract

To understand the genetic variation underlying atrial fibrillation (AF), the most common cardiac arrhythmia, we performed a genome-wide association study (GWAS) of > 1 million people, including 60,620 AF cases and 970,216 controls. We identified 163 independent risk variants at 111 loci and prioritized 165 candidate genes likely to be involved in AF. Many of the identified risk variants fall near genes where more deleterious mutations have been reported to cause serious heart defects in humans or mice *(MYH6, NKX2-5, PITX2, TBC1D32*, TBX5),^1,2^ or near genes important for striated muscle function and integrity (e.g. *MYH7, PKP2, SSPN, SGCA)*. Experiments in rabbits with heart failure and left atrial dilation identified a heterogeneous distributed molecular switch from *MYH6* to *MYH7* in the left atrium, which resulted in contractile and functional heterogeneity and may predispose to initiation and maintenance of atrial arrhythmia.

## Results

We tested the association between 34,740,186 genetic variants (minor allele frequency [MAF] > 2.5×10^−5^) and AF, comparing a total of 60,620 cases and 970,216 controls of European ancestry from 6 contributing studies (HUNT, deCODE, MGI, DiscovEHR, UK Biobank, and the AFGen Consortium) (**Supplementary Table 1**). We identified 111 genomic regions with at least 1 genetic variant associated with AF (P-value < 5 × 10^−8^). Of these, 80 loci have not previously been reported (**Supplementary Figure 1**, **Figure 1**, **Supplementary Table 2**, and **Supplementary Table 3**). Based on approximate stepwise conditional analyses,^3^ we identified 52 additional genetic risk variants within the 111 loci that demonstrated genome-wide statistically significant association with AF (**Supplementary Table 4**) that were independent of the locus index variant (LD r^2^ < 0.05). We applied the widely used GWAS P-value significance threshold of P-value < 5×10^−8^. Some have suggested to use a more stringent threshold of 5×10^−9^ when testing millions of imputed markers.^4^ If we had applied this threshold, we would still identify 94 loci, 63 of which have not been previously reported (**Supplementary Table 2**).

**Figure 1.**
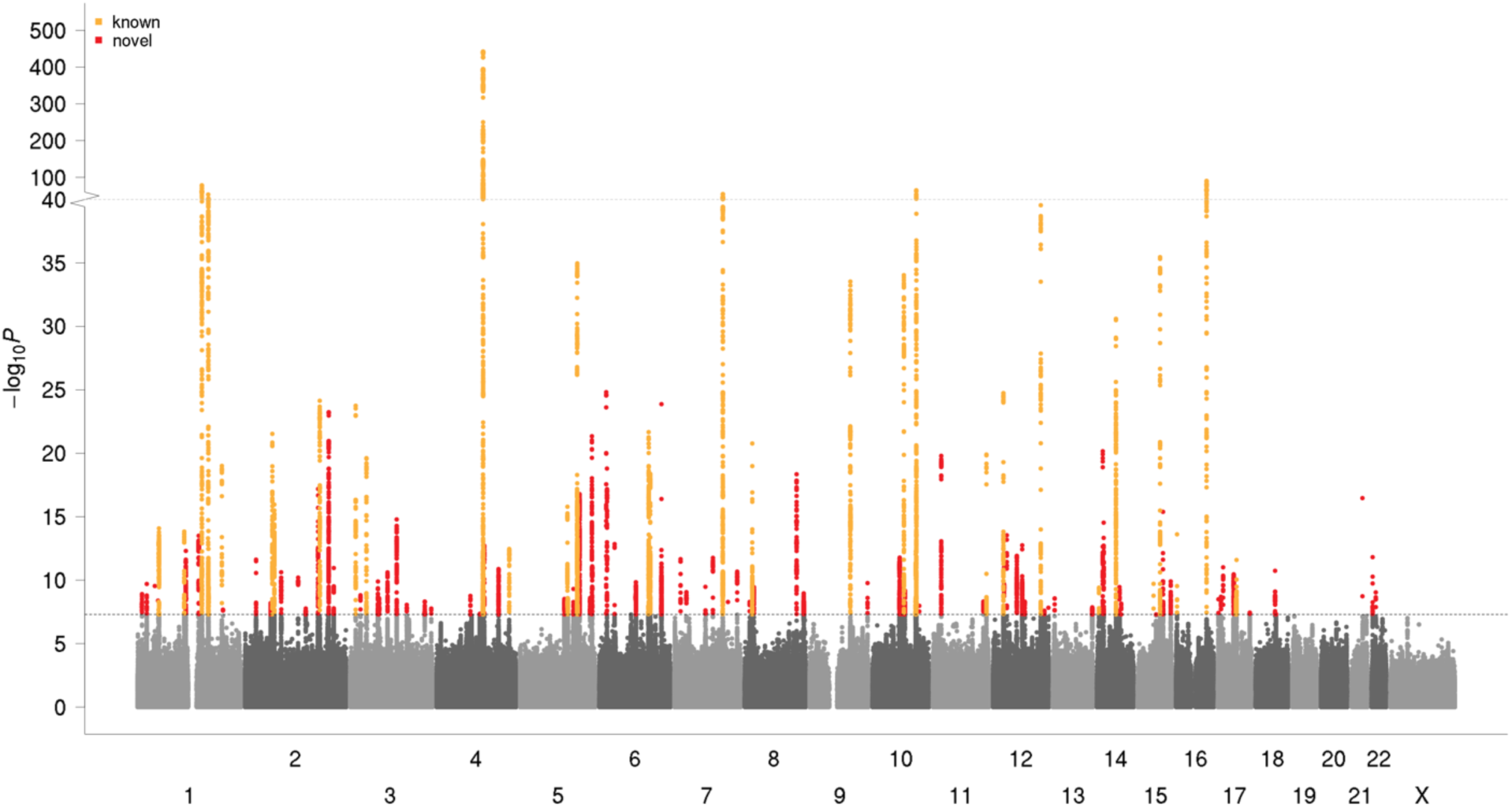
Manhattan plot showing known (orange) and novel (red) loci associated with atrial fibrillation. The x-axis represents the genome in physical order whereas the y-axis represents P-values (−log_10_[P-value]) of association.

Of the 35 loci previously reported for AF (**Supplementary Table 3**), we identified genome-wide significant association (P-value < 5×10^−8^) at 31 (89%) after excluding results from the previously published AFGen Consortium, which has published the majority loci reported to date (**Supplementary Table 5**).^5^ The 4 loci not captured comprised 3 loci discovered in East Asian populations *(KCNIP1, NEBL*, and *CUX2)* and 1 locus *(PLEC)* for which we did not have data on the previously reported missense variant.^6^ To further test the validity of our findings, we performed a heterogeneity test for the 111 index variants across the 6 contributing studies. Of the 111 index variants, only 2 index variants showed evidence for heterogeneity in the effect size across the 6 contributing studies (P-value < 0.05/111 = 4.5×10^−4^) (**Supplementary Table 2**). Both of these index variants represent loci that have previously been established as associated with AF across multiple studies (near *PRRX1, PITX2)* (**Supplementary Table 3**). These findings demonstrate a high external validity of our results.

To understand the biology underlying the 111 AF-associated loci, we employed a number of approaches, including ‘Data-driven Expression Prioritized Integration for Complex Traits’ (DEPICT)^7^ to identify cell types and tissues in which genes at AF-associated variants are likely to be preferentially expressed. Based on 37,427 human microarray expression samples from 209 different tissues and cell types, we observed a statistically significant enrichment for atrial (P-value = 2.4×10^−5^), atrial appendage (P-value = 2.8×10^−5^), heart (P-value = 5.2×10^−5^), and ventricular tissues (P-value = 1.1×10^−4^) (Figure 2a and **Supplementary Table 6**). We further applied DEPICT to detect gene sets that were enriched for genes at AF-associated loci. Of the 14,461 gene sets we tested, 889 were enriched (false discovery rate [FDR] < 0.05) for genes at AF-associated loci (Figure 2b and **Supplementary Table 7**). The highlighted gene sets in general point to biological processes related to cardiac development and morphology along with structural remodeling of the myocardium. These findings are in line with recent reports which have linked AF with rare coding variants in the sarcomere genes *MYH6* and *MYL4* and in the multidomain cyto-skeletal linking protein *PLEC* along with more common coding variants in *TTN*, essential for the passive elasticity of heart and skeletal muscle.^8,9,6,10^

**Figure 2.**
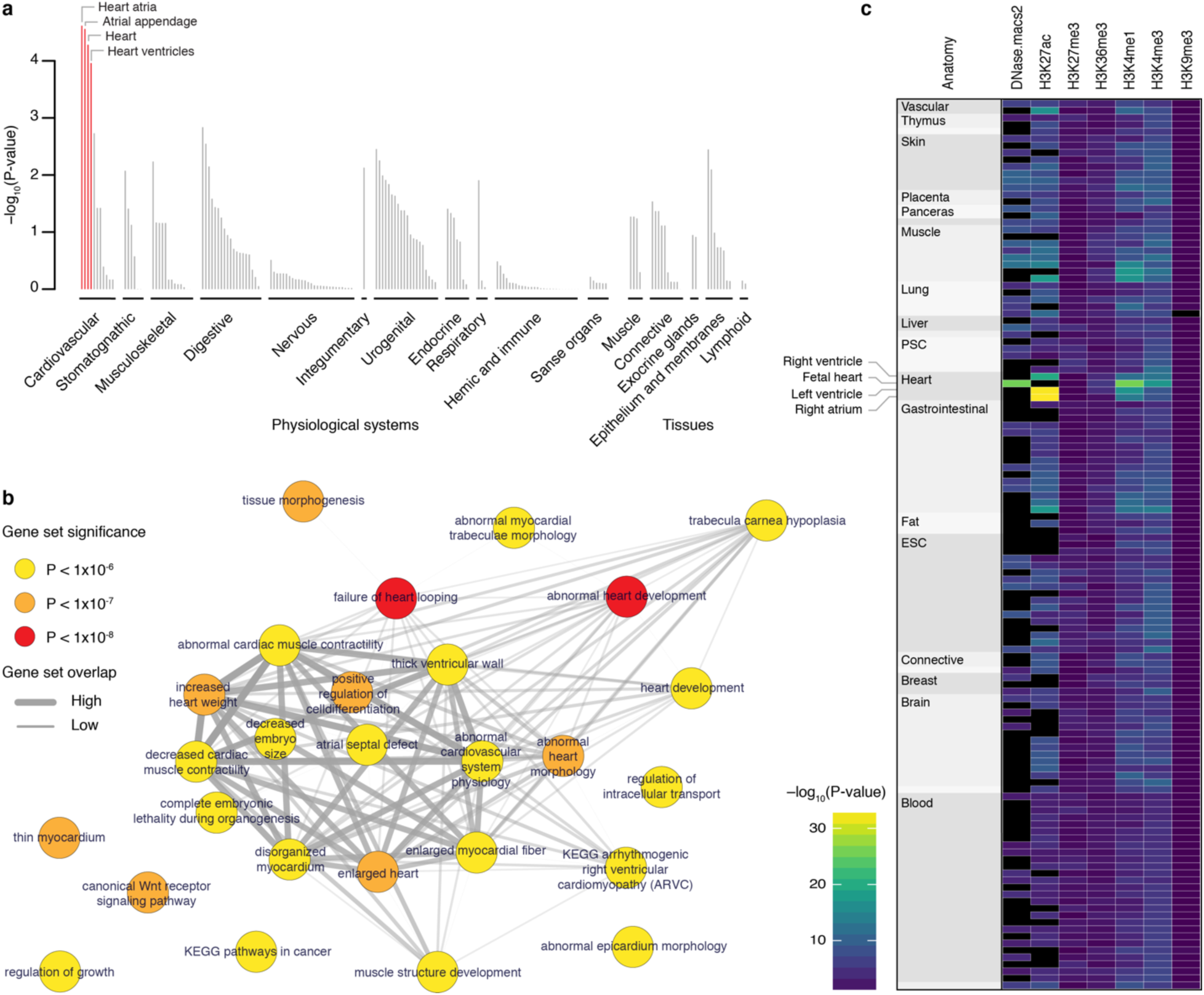
Tissues, reconstituted gene sets, and regulatory elements implicated in atrial fibrillation. **a)** Based on expression patterns across 37,427 human mRNA microarrays, DEPICT predicted genes within atrial fibrillation-associated loci to be highly expressed across various cardiac tissues. Tissues are grouped by type and significance. Red columns represent statistically significant tissues following Bonferroni correction (P-value < 0.0002). **b)** Top (P < 1×10^−6^) reconstituted gene sets (out of 826 with FDR < 0.05 and after exclusion of ‘gene subnetworks’) found by DEPICT to be significantly enriched by genes in atrial fibrillation-associated loci. Each node, colored according to the permutation P-value, represents a gene set and the grey connecting lines represent pairwise overlap of genes within the gene sets. **c)** Heatmap indicating the overlap between fibrillation–associated risk variants and regulatory elements across 127 Roadmap Epigenomics tissues (each represented by a row) using GREGOR. Black indicates no data.

Although we could identify protein-altering variants at n = 21 loci, comprising either the index variant (n = 2 loci) or a variant in high linkage disequilibrium (LD) (r^2^) with the index variant (n = 19 loci; **Supplementary Table 8**), we noted that most associated risk variants are in the non-coding genome (159 of 163 independent risk variants). To assess the potential function of associated non-coding variants, we tested for enrichment of AF-associated variants with a variety of regulatory features including DNase I hypersensitive sites (DHS), histone methylation marks, transcription factor binding sites, and chromatin states in a variety of cell and tissue types available from Roadmap Epigenomics^11^ using ‘Genomic Regulatory Elements and Gwas Overlap algoRithm’ (GREGOR).^12^ This method tests if the number of AF-associated index variants, or their LD proxies, overlap with the corresponding regulatory feature more often than expected when compared to a permuted control sets. Of 787 combinations of regulatory features and tissues examined (**Supplementary Table 9**), we found that AF-associated variants were most strongly associated with: active enhancers as indicated by H3K27ac in right atrium (P-value = 2×10" ^33^; 2.9x enrichment); H3K27ac in left ventricle (P-value = 3×10^−33^; 2.6x enrichment); and in fetal heart tissue we found strong enrichment with H3K4me1 (P-value = 9×10^−27^; 2.0x enrichment) and open chromatin (P-value = 2×10^−26^; 2.1x enrichment) (**Figure 2c**, **Supplementary Figure 2** and **Supplementary Table 9**). This suggests that some loci are important in transcriptional regulation in the adult heart, in development of the fetal heart, or both.

To further enhance the biological understanding of the AF-associated loci, we identified candidate functional genes. There were 3,072 genes or transcripts for which the transcription region overlapped (see Methods) at least one variant in the 111 loci. We prioritized biological candidate genes which: i) harbored a protein-altering variant that was in high LD (r^2^ > 0.80; **Supplementary Table 8**) or was itself the locus index variant; ii) expression levels were associated and colocalized with AF-associated variants (P-value < 1.14 × 10^−9^ in GTEx consortium data);^13^ iii) were highlighted by DEPICT (FDR < 0.05); or iv) were nearest to the index variant in a locus. Using these criteria, we prioritized 165 target genes (**Supplementary Table 2**, **Supplementary Table 10**, and **Supplementary Table 11**).

To identify tissues in which the 165 prioritized candidate genes showed enhanced expression, we used ‘Tissue Specific Expression Analysis’ (TSEA)^14^ and found enrichment in heart (P-value = 5×10^−12^), muscle (P-value = 1×10^−9^) and blood vessel tissues (P-value = 2×10^−9^). To assess the empirical significance of these results, we performed 1,000 permutations of the same number of genes selected: i) randomly from the genome and ii) subsets of the 3,072 genes within the 111 AF loci. We determined that the observed P-values were substantially more significant than expected by chance (**Figure 3**). These findings support that the genes we prioritized are strong candidates for being involved in AF.

**Figure 3.**
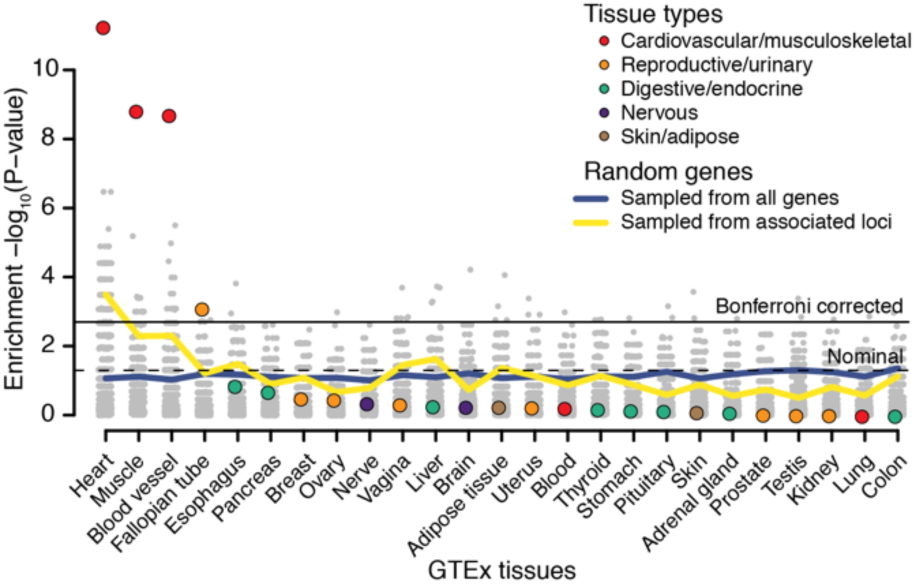
Significance of the expression enrichment for the candidate genes. This figure compares the tissue-specific gene expression enrichment for the 165 biological candidate genes (colored dots) to a null distribution derived by randomly selecting same number of genes from the whole genome or from the associated loci. The grey dots are the P-values for each of the permutations for the randomized tests (1,000 for both sampling scenarios for each tissue) and the blue and yellow lines represent the per-tissue P-value thresholds comparable to a false positive rate of 0.05.

Interestingly, we identified as functional candidates at least 20 genes likely to be involved in cardiac and skeletal muscle function and integrity *(AKAP6, COL25A, CFL2, DPT, MYH6, MYH7, MYO18B, MYO1C, MYOCD, MYOT, MYOZ1, MYPN, PKP2, RBM20, SGCA, SSPN, SYNPO2L, TTN, TTN-AS, WIPF1);* these included SGCA and *SSPN*, which have been associated with muscular dystrophies,^15,16^ and *PKP2* which has been associated with arrhythmogenic right ventricular cardiomyopathy.^17^ We also identified at least 13 genes likely to be involved in mediation of developmental events *(EPHA3, GTF2I, HAND2, MYH6, NAV2, NKX2-5, PITX2, SLIT3, SOX15, SOX5, TBC1D32, TBX5, TGFB3)* along with genes likely to be involved in intracellular calcium handling in the heart *(CALU, CAMK2D, CASQ2, PLN, S100A7A)*, angiogenesis *(TNFSF12, TNFSF12-TNFSF13)*, hormone signaling (ESR2, IGF1R, JMJD1C, NR3C1, THRB1), and function of cardiac ion channels *(GRIK4, KCNC2, KCND3, KCNH2, KCNJ5, KCNN2, KCNN3, SCN10A, SCN5A, SLC9B1)*.

We tested the 111 AF index variants for association with 123 electrocardiogram (ECG) parameters in 62,974 Icelanders in sinus rhythm, after exclusion of AF cases (**Supplementary Figure 3**). Sixty variants were associated with at least one ECG parameter when we controlled for a false discovery rate of 0.05 at the variant level, 39 of which were novel AF variants including many with substantial ECG effects, such as the variants near *NACA, THRB, CAMK2D, NKX2-5*, and *CDKN1A*.

For the locus around index variant rs422068 on chromosome 14, our approach prioritized *MYH6* and *MYH7* as the most likely functional genes (**Supplementary Table 2**). *MYH6* encodes myosin heavy chain alpha (a-MyHC), a major component of the thick filaments of the *contractile apparatus* in adult atria, and hence important for atrial contraction.^18^ *MYH7* encodes β-MyHC, a slower acting isoform,^19^ and is mainly expressed in the ventricles of the human heart. It has been established that *MYH6* and *MYH7* are regulated in an inverse manner, and that in heart failure and other cardiac disorders in humans, β-MHC is upregulated, whereas α-MHC is downregulated, resulting in diminution of cardiac performance.^20^ Whether these changes occur also in the atria has not previously been addressed.

To explore potential mechanisms of *MYH6* and *MYH7* in AF, we developed an ischemic heart failure model for AF in rabbits. Ischemia was produced by chronic ligation of the left circumflex artery (LCX) during thoracotomy with subsequent development of ischemic heart failure (> 4 weeks post operatively) and profound left atrial dilation. We found that *MYH7* expression was only detectable in the heart failure remodeled left atrium (**Figure 4**). The control left atrium did not express detectable levels of *MYH7* and exclusively expressed *MYH6*. More importantly, in the dilated left atrium, *MYH7* expression was heterogeneously distributed and thus resulted in contractile heterogeneity, which may have predisposed hearts to develop long-lasting AF, particularly when intra-atrial pressure was increased to 10cm H_2_O. Control hearts did not develop long-lasting AF until intra-atrial pressure was increased to 30cm H_2_O. (**Figure 4**, **Supplementary Figure 4**). Altogether, this experiment demonstrated that a *MYH6* to *MYH7* switch in the atria may accompany or predispose to atrial fibrillation, and that the expression of both the faster and slower myosin heavy chain forms may predispose to arrhythmia through contractile heterogeneity.

**Figure 4.**
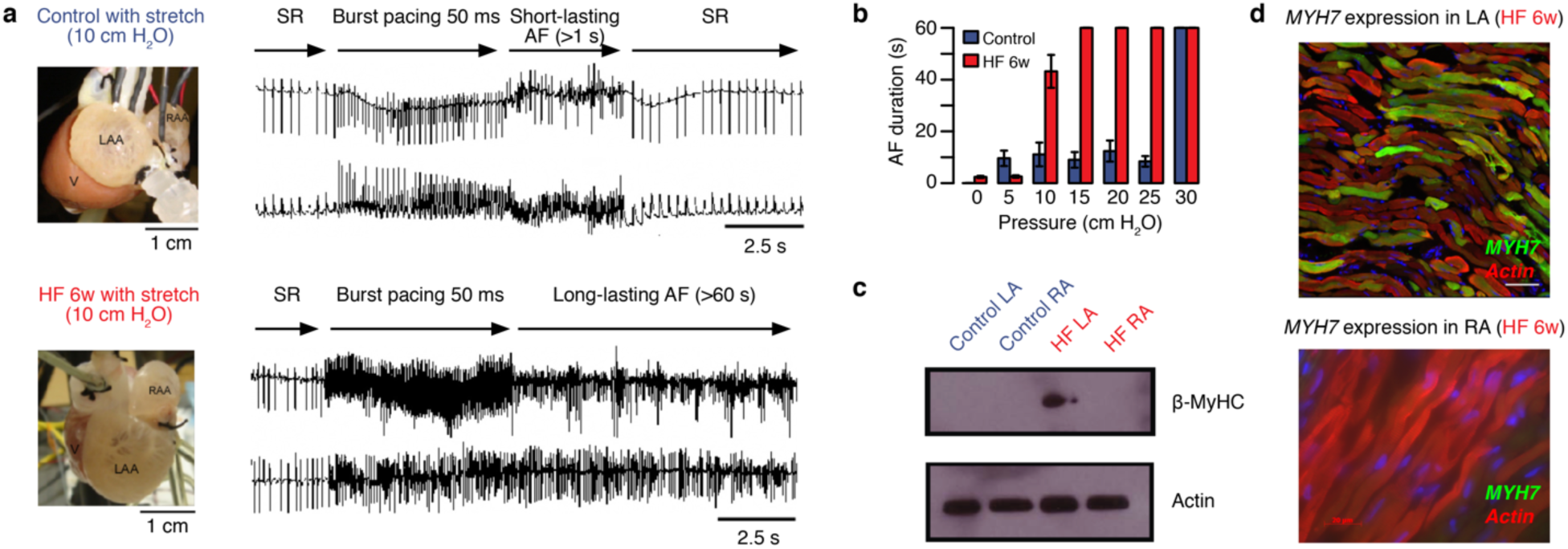
Atrial fibrillation (AF) is associated with heterogeneous changes in left atrial myosin isoform expression. **a)** Langendorff-perfused rabbit hearts from control (blue, top) or heart failure (HF) rabbits (red, bottom panel) were tested for AF-inducibility and duration following burst pacing at 50ms cycle length. HF was induced by chronic left circumflex artery ligation and was allowed to develop over 6 weeks. During HF progression, severe left atrial hypertrophy occurred. **b)** HF hearts developed long lasting AF (> 60s) when intra-atrial pressure was increased to 10 cm H_2_O. On the other hand, control hearts did not develop long lasting AF until intra-atrial pressure was increased to 30cm H_2_O. **c)** Western blotting for MYH7 gene expression (β-MyHC protein) indicates MYH7 expression exclusively in the remodeled HF left atrium. **d)** Immunostaining and confocal microscopy revealed heterogeneous MYH7 gene expression (green) in the HF left atrium. Consistent with Western blotting data, the HF right atrium (RA) did not express MYH7.

Next, we investigated whether any of the 165 biological candidate genes that we identified could potentially represent a novel drug target for already developed drugs or drugs undergoing development by querying the Drug-Gene Interaction Database.^21^ We found one or more potential drug or substance-interactions for 39 of the 165 prioritized genes, totaling 523 drugs. Of these, 77 drugs targeting 16 genes are already known to be able to control or trigger AF or other cardiac arrhythmias (**Supplementary Table 12**). Gene-drug interactions worth highlighting include the interaction between *MYH6* and *MYH7* and omecamtiv mecarbil and the interaction between *KCNH2* and rottlerin. Omecamtiv mecarbil is a cardiac-specific myosin activator which is currently being tested for treatment of heart failure^22^. Rottlerin, a natural product isolated from the tree Mallotus philippensis, has been shown to increase cardiac contractile performance and coronary perfusion through mitochondrial BK_Ca++_ channel activation in rat hearts.^23^ Whether these or the other highlighted drugs can impact AF needs further evaluation but the findings can be used as a foundation for directing future functional experiments and clinical trials.

Finally, we constructed polygenic risk scores using weighted effect estimates generated from the deCODE sample (13,471 AF cases vs. 358,161 controls). We tested the performance of the deCODE-based weighted polygenic risk score against prevalent AF in the Norwegian HUNT study (6,337 cases vs. 61,607 controls) using a variety of different thresholds of association P-values and LD pruning thresholds. We observed the highest area under the receiver operating curve using genotype dosages for markers with a P-value < 5×10^−5^ that were pruned using an LD r^2^-threshold of 0.8 (n = 725 risk markers; AUC = 57.7%, **Supplementary Figure 5**). We used this optimized polygenic risk score to test for association with 1,494 International Classification of Diseases (ICD) code-defined disease groups in UK Biobank participants of white British ancestry.^24^ In addition to a strong association with AF (P-value = 7×10^−374^), we found association to 33 mainly cardiovascular conditions (P-value < 0.05/1,494 = 3.3×10^−5^), including palpitations, mitral valve disorders, hypertension, heart failure, ischemic heart disease, and stroke (**Supplementary Table 13** and **Supplementary Figure 6**). However, when participants diagnosed with any type of cardiac arrhythmia (n = 24,681) were excluded from the analyses to avoid assessment bias, the AF risk score was not associated with any ICD disease group (P-value > 3.3×10^−5^). This suggests that the score is specific for AF or cardiac arrhythmia and that the additional associations that we identified were mediated through AF, either as a result of a more thorough clinical examination (e.g valvular disease) or because AF is a likely intermediate step towards the disease (e.g. stroke).

In summary, we substantially increased the number of genome-wide significant risk variants for AF through a large GWAS meta-analysis. Based on pathway and functional enrichment analyses along with prioritization of functional candidate genes we anticipate that many AF risk variants act in the developing heart or impact AF via structural remodeling of the myocardium in the form of an ‘atrial cardiomyopathy’^25^ as a response to atrial stress in the adult heart. This finding needs confirmation but provides a strong foundation for directing future functional experiments to better understand the biology underlying AF.

## Methods

### Discovery cohorts

More details on some cohorts are provided in the Supplementary Appendix. **HUNT**: The Nord-Trøndelag Health Study (HUNT) is a population-based health survey conducted in the county of Nord-Trøndelag, Norway, since 1984.^26^ We used a combination of hospital, out-patient, and emergency room discharge diagnoses (ICD-9 and ICD-10) to identify 6,337 AF cases and 61,607 AF-free controls with genotype data. **DeCODE**: The Icelandic AF population consisted of all patients diagnosed with AF (International Classification of Diseases (ICD) 10 code I.48 and ICD 9 code 427.3) at Landspitali, The National University Hospital, in Reykjavik, and Akureyri Hospital (the two largest hospitals in Iceland) from 1987 to 2015. All AF cases, a total of 13,471, were included. Controls were 358,161 Icelanders recruited through different genetic research projects at deCODE genetics. Individuals in the AF cohort were excluded from the control group. **MGI**: The Michigan Genomics Initiative (MGI) is a hospital-based cohort collected at Michigan Medicine, USA. Atrial fibrillation cases (n = 1,226) were defined as patients with ICD-9 billing code 427.31 and controls were individual without AF, atrial flutter, or related phenotyps (ICD-9 426-427.99). **DiscovEHR**: The DiscovEHR collaboration cohort is a hospital-based cohort including 58,124 genotyped individuals of European ancestry from the ongoing MyCode Community Health Initiative of the Geisinger Health System, USA. AF cases (n = 6,679) were defined as DiscovEHR participants with at least one electronic health record problem list entry or at least two diagnosis code entries for two separate clinical encounters on separate calendar days for ICD-10 I48: atrial fibrillation and flutter. Corresponding controls (n = 41,803) were defined as individuals with no electronic health record diagnosis code entries (problem list or encounter codes) for ICD-10 I48. **UK biobank**: The UK Biobank is an population-based cohort collected from multiple sites across the United Kingdom.^24^ Cases of AF were selected using ICD-9 and ICD-10 codes for AF or atrial flutter (ICD-9 427.3 and ICD-10 I48). Controls were participants without any ICD-9 or ICD-10 coded specific for AF, atrial flutter, other cardiac arrhythmias, or conduction disorders. **AFGen Consortium**: Published AF association summary statistics from 31 cohorts representing 17,931 AF cases and 115,142 controls were obtained from the authors.^5^

### Genotyping array, imputation and association analysis

**HUNT**: Genotyping was performed at the Norwegian University of Science and Technology (NTNU) using the Illumina HumanCore Exome v1.0 and v1.1. Quality control was performed at the marker and sample level. A total of 2,201 individuals were whole genome sequenced at low-pass and genotype calls were generated using gotCloud pipeline (https://genome.sph.umich.edu/wiki/GotCloud). Variants from the HUNT low-pass genomes were imputed into HRC samples and vice-versa to generate a single imputation reference panel of ^~^34,000 individuals including 2,201 study-specific samples. Imputation was performed using Minimac3 and variants with imputation r^2^ > 0.3 were take forward. We performed testing for association with AF using a generalized mixed model including covariates birth year, sex, genotype batch, and principal components (PC) 1–4 as implemented in SAIGE.^27^ **DeCODE**: The study is based on whole-genome sequence data from 15,220 Icelanders participating in various disease projects at deCODE genetics. The sequencing was done using Illumina standard TruSeq methodology to a mean depth of 35x (SD 8).^8^ Autosomal SNPs and INDEL’s were identified using the Genome Analysis Toolkit version 3.4.0.^28^ Variants that did not pass quality control were excluded from the analysis according to GATK best practices. Genotypes of the sequence variants identified through sequencing (SNPs and indels) were then imputed into 151,677 Icelanders chip typed using Illumina SNP chips and their close relatives (familial imputation).^29^ Variants for the meta-analysis were selected based on matching with either the 1000g reference panel (Phase 3) or the Haplotype Consortium reference panel^30^ (based on allele, frequency and correlation matching). Logistic regression was used to test for association between SNPs and AF, treating disease status as the response and allele counts from direct genotyping or expected genotype counts from imputation as covariates. Other available individual characteristics that correlate with phenotype status were also included in the model as nuisance variables. These characteristics were: sex, county of birth, current age or age at death (first and second order terms included), blood sample availability for the individual and an indicator function for the overlap of the lifetime of the individual with the time span of phenotype collection. To account for inflation in test statistics due to cryptic relatedness and stratification, we applied the method of linkage disequilibrium (LD) score regression.^31^ The estimated correction factor for AF based on LD score regression was 1.38 for the additive model. **MGI**: Genotyping was performed at the University of Michigan using the Illumina Human Core Exome v1.0 and v1.1. Quality control was performed at the marker and sample level. Imputation of variants from the HRC reference panel was performed using the Michigan Imputation Server (https://imputationserver.sph.umich.edu/index.html) and variants with imputation r^2^ > 0.3 were included. Association with AF was determined using the Firth bias-corrected logistic likelihood ratio test^32^ with adjustment for age, sex, and PC1–4. **DiscovEHR**: Aliquots of DNA were sent to Illumina for genotyping on the Human OmniExpress Exome Beadchip. All individuals of European ancestry, as determined using PC analysis, were imputed to the HRC Reference Panel using the Michigan Imputation Server. Markers with imputation r^2^ > 0.3 and MAF > 0.001 were carried forward for analysis. BOLT-LMM^33^ was used to analyze BGEN dosage files, and variants were tested for association with atrial fibrillation under an additive genetic model, adjusting for sex, age, age^2^, and the first four PCs of ancestry; additionally, a genetic relatedness matrix (calculated using variants with MAF > 0.001, per-genotype missing data rate < 1%, and Hardy–Weinberg equilibrium P-value < 10^−15^) was included as a random-effects variable in the model.^34^ **UK biobank**: Details on quality control, genotyping and imputation can be found elsewhere.^35^ In brief, study participants were genotyped using two very similar genotyping arrays (Applied Biosystems™ UK BiLEVE Axiom™ Array and UK BioBank Axiom™ Array) designed specifically for the UK Biobank. Phasing and imputation was done by the UK Biobank analyses team based on the HRC reference panel and the UK10K haplotype resource.^35^ We restricted our analyses to HRC-imputed markers only as there have been reports of incorrect estimates for non-HRC markers in the first 500,000 people release from UK Biobank. We performed testing for association with AF in people of white British ancestry using a generalized mixed model including covariates birth year, sex, genotype batch, and principal PC 1–4 as implemented SAIGE.^27^

### Meta-analysis

We included all markers that were available for analyses in any of the 6 contributing studies. For the DiscoverEHR that applied the BOLT-LMM mixed model, we obtained an approximation of the allelic log-OR and corresponding variance from the linear model as described previously.^36^ Following this, we performed a meta-analyses using the inverse variance method implemented in the software package METAL (http://genome.sph.umich.edu/wiki/METAL_Documentation).^37^ When estimating the cross-cohort allele frequencies, we only included participating studies where individuals were sampled independent of AF status (HUNT, deCODE, MGI, DiscoverEHR, UK Biobank). This was done to avoid sampling bias. Heterogeneity tests were performed as implemented in METAL.^37^

### Definition of independent loci

Independent loci were defined as genetic markers > 1Mb and > 0.25 cM apart in physical and genomic distance, respectively, with at least 1 genetic variant associated with AF at a genome-wide significance threshold of P-value < 5 × 10^−8^. The lower loci boarders were defined as the genome-wide statistically significant marker within the loci with the lowest genomic position minus 1Mb. The upper loci boarders were defined as the genome-wide statistically significant marker within the loci with the highest genomic position plus 1Mb.

### Linkage disequilibrium (LD) estimation

We used 5,000 unrelated individuals that were randomly sampled among the HUNT Study participants to calculate calculated LD r^2^ using the software PLINK1.9 (https://www.cog-genomics.org/plink/1.).

### Approximate, stepwise conditional analyses

To identify independent risk variants within the identified AF-associated loci, we used the COJO-GCTA software (http://cnsgenomics.com/software/gcta/) to performed approximate, stepwise conditional analyses based on summary statistics from the meta-analyses and a LD-matrix obtained from 5,000 unrelated individuals randomly sampled from the HUNT Study.^3^ Only variants with MAF > 0.01 were included in the analyses and variants were only considered truly independent if they were not in LD (r^2^ < 0.05) with the locus index variant and any of the other independent risk variants.

### Identifying candidate functional genes using DEPICT

We employed DEPICT (https://data.broadinstitute.org/mpg/depict/) to identify 1) the most likely causal gene at associated loci, 2) reconstituted gene sets enriched for AF loci, and 3) tissues and cell types in which genes that form associated loci are highly expressed.^7^ DEPICT uses gene expression data derived from a panel of 77,840 mRNA expression arrays^38^ together with 14,461 existing gene sets defined based on molecular pathways derived from experimentally verified protein-protein interactions,^39^ genotype-phenotype relationships from the Mouse Genetics Initiative,^40^ Reactome pathways,^41^ KEGG pathways,^42^ and Gene Ontology (GO) terms.^43^ Based on similarities across the microarray expression data, DEPICT reconstitutes the 14,461 existing gene sets by assigning each gene in the genome a likelihood of membership in each gene set. Using these precomputed gene sets and a set of trait-associated loci, DEPICT quantifies whether any of the 14,461 reconstituted gene sets are significantly enriched for genes in the associated loci and prioritizes genes that share predicted functions with genes from the other associated loci more often than expected by chance. Additionally, DEPICT uses a set of 37,427 human mRNA microarrays to identify tissues and cell types in which genes from associated loci are highly expressed (all genes residing within a LD of r^2^ > 0.5 from index variant).

We ran DEPICT using all AF-associated index variants and variants identified through stepwise conditional analyses. For the gene sets significantly enriched for AF-associated loci (P-value < 1 × 10^−6^, FDR <0.05), we computed a weighted pairwise similarity based on the number of overlapping genes for genes with a Z score < 4.75 (corresponding to P-value < 1 × 10^−6^) for being part of the gene set. For gene sets with no genes with a Z score < 4.75, we included the 3 most significant genes as done previously.^44^

### GREGOR

We tested for enrichment of index variants with functional domains using the software GREGOR (http://csg.sph.umich.edu/GREGOR/).^12^ This method tests for an increase in the number of AF-associated index variants, or their LD proxies, overlapping with the regulatory feature more often than expected by chance by comparing to permuted control sets where the index variant is matched for frequency, number of LD proxies and distance to the nearest gene. We use a saddle-point approximation to estimate the P-value by comparing to the distribution of permuted statistics.^12^ We ran GREGOR using all AF-associated index variants along with variants identified through stepwise conditional analyses.

### Identification of expression quantitative trait loci (eQTLs) using GTEx data

We performed eQTL look-up using the GTEx database (http://gtexportal.org)^13^ version 6p, which holds cis-eQTLs expression data of up to 190 million single nucleotide variants across 44 tissues, by searching for all AF-associated loci index variants, all independent risk variants identified from the stepwise conditional analyses, and any variants in strong LD (r^2^ > 0.80) with these variants using an eQTL significance threshold of P < 1.14 × 10^−9^ (5 × 10^−8^ / 44 tissues). For all statistically significant genes, we queried all markers in the GTEx database that affected the expression of the affected genes and tested if the eQTLs markers colocalized with the GWAS signal as described previously.^45^

### Ischemic heart failure model of atrial fibrillation susceptibility

Ischemic heart failure was modeled using a previously described rabbit model of left circumflex artery ligation. In this model, the left atrium progressively dilates following the ischemic insult as heart failure develops. Figure 4a shows images of Langendorff perfused hearts of control and heart failure (HF) animals highlighting the overt dilation of the left atrium in HF. With equivalent left atrial pressure (10 cm H_2_O) AF was induced in each condition with high frequency burst pacing as shown in the ECG traces and done before.^46^ Protein expression analysis were performed using western blot.

### Tissue Specific Expression Analysis (TSEA)

The TSEA analyses were performed using the R software pSI package

(http://genetics.wustl.edu/jdlab/psi_package/).^14^ For the calculations, pre-defined pSI values provided by the pSI package creators were used. To get null distributions for the P-values for the prioritized genes, we performed two sets of permutations; randomly selected from the entire human genome and randomly selected from the associated loci (also matching the number of genes picked in each of the loci). In both scenarios one thousand permutations were done.

### Electrocardiogram data

ECG data was collected from Landspitali University Hospital in Reykjavik and included all ECGs obtained and digitally stored from 1998 to 2015, including a total of 434,000 ECGs from 88,217 individuals. A total of 289,297 ECGs of 62,974 individuals were sinus rhythm (heart rate 50–100 beats per minute) ECGs of individuals without the diagnosis of AF. The ECGs were digitally recorded with the Philips PageWriter Trim III, PageWriter 200, Philips Page Writer 50 and Phillips Page Writer 70 cardiographs and stored in the Philips TraceMasterVue ECG Management System. These were ECGs obtained in all hospital departments, from both inpatients and outpatients. Digitally measured ECG waveforms and parameters were extracted from the database for analysis. The Philips PageWriter Trim III QT interval measurement algorithm has been previously described and shown to fulfill industrial ECG measurement accuracy standards.^47^ The Philips PR interval and QRS complex measurements have been shown to fulfill industrial accuracy standards.^48^

We tested 111 genome-wide significant and replicated AF variants for association with 123 ECG measurements using a linear mixed effects model implemented in the Bolt software package,^33^ treating the ECG measurement as the response and the genotype as the covariate. All measures except heart rate and QT corrected are presented for all 12 ECG leads. For this analysis, we used 289,297 sinus rhythm ECGs (heart rate 50–100 beats per minute) from 62,974 individuals who have not been diagnosed with AF according to our databases. This was done to assess the effect of the AF variants on ECG measures and cardiac electrical function in the absence of AF. Individuals with pacemakers were also excluded. The ECG measurements were adjusted for sex, year of birth, and age at measurement and were subsequently quantile standardized to have a normal distribution. For individuals with multiple ECG measurements, the mean standardized value was used. We assume that the quantitative measurements follow a normal distribution with a mean that depends linearly on the expected allele at the variant and a variance-covariance matrix proportional to the kinship matrix.^49^ Since 123 traits were tested, the Benjamini-Hochberg FDR procedure controlling the FDR at 0.05 at each marker was used to account for multiple testing.

### Polygenic risk score

Using dosage-weighted effect estimates obtained from the Iceland-based deCODE population, we constructed 20 GWAS-based polygenic risk cores by combining genetic markers across different GWAS P-value thresholds (P-value < 5 × 10^−4^, P-value < 5 × 10^−5^, P-value < 5 × 10^−6^, P-value < 5 × 10^−7^, P-value < 5 × 10^−8^) and LD cut-offs (r^2^ < 0.2, r^2^ < 0.4, r^2^ < 0.6, r^2^ < 0.8). We evaluated the performance of each of the 20 polygenic risk scores against AUC for predicting prevalent AF in the Norwegian-specific HUNT Study using a logistic regression.

### Phenome-wide association analyses

We used a previously published scheme to defined disease-specific binary phenotypes by combining hospital ICD-9 codes into hierarchical PheCodes, each representing a more or less specific disease group.^50^ ICD-10 codes were mapped to PheCodes using a combination of available maps through the Unified Medical Language System(https://www.nlm.nih.gov/research/umls/) and other sources, string matching, and manual review. Study participants were labeled a PheCode if they had one or more of the PheCode-specific ICD codes. Cases were all study participants with the PheCode of interest and controls were all study participants without the PheCode of interest or any related PheCodes. Gender checks were performed, so PheCodes specific for one gender could not mistakenly be assigned to the other gender. The association between the optimized polygenic risk score and each of the defined phenotypes where tested using a logistic regression adjusted for sex and birth year.

## Acknowledgements and Funding

The Nord-Trøndelag Health Study (The HUNT Study) is a collaboration between HUNT Research Centre (Faculty of Medicine, NTNU, Norwegian University of Science and Technology), Nord-Trøndelag County Council, Central Norway Health Authority, and the Norwegian Institute of Public Health. J.B.N. was supported by grants from the Danish Heart Foundation and the Lundbeck Foundation. T.J.H was supported by an American Heart Association Scientist Development Grant (0735464Z). J.A.S. was supported by National Institute of Health (NIH) grant R01-HL124232. The K.G. Jebesen center for genetic epidemiology is financed by Stiftelsen Kristian Gerhard Jebsen, Faculty of Medicine and Health Sciences Norwegian University of Science and Technology (NTNU) and Central Norway Regional Health Authority.

